# Structural elements required for coupling ion and substrate transport in the neurotransmitter transporter homolog LeuT

**DOI:** 10.1101/283176

**Authors:** Yuan-Wei Zhang, Sotiria Tavoulari, Steffen Sinning, Antoniya A. Aleksandrova, Lucy R. Forrest, Gary Rudnick

**Author notes:** Corresponding Author: Gary Rudnick, Department of Pharmacology, Yale University School of Medicine, 333 Cedar Street, New Haven, CT 06520-8066.

## Abstract

The coupled transport of ions and substrates allows transporters to accumulate substrates using the energy of transmembrane ion gradients and electrical potentials. During transport, conformational changes that switch accessibility of substrate and ion binding sites from one side of the membrane to the other must be controlled so as to prevent uncoupled movement of ions or substrates. In the Neurotransmitter:Sodium Symporter (NSS) family, Na^+^ stabilizes the transporter in an outward-open state, thus decreasing the likelihood of uncoupled Na^+^ transport. Substrate binding, in a step essential for coupled transport, must overcome the effect of Na+, allowing intracellular substrate and Na^+^ release from an inward-open state. However, the specific elements of the protein that mediate this conformational response to substrate binding are unknown. Previously, we showed that in the prokaryotic NSS transporter LeuT, the effect of Na^+^ on conformation requires the Na2 site, where it influences conformation by fostering interaction between two domains of the protein (JBC 291: 1456, 2016). Here, we used cysteine accessibility to measure conformational changes of LeuT in *E. coli* membranes. We identified a conserved tyrosine residue in the substrate binding site required for substrate to convert LeuT to inward-open states by establishing an interaction between the two transporter domains. We further identify additional required interactions between the two transporter domains in the extracellular pathway. Together with our previous work on the conformational effect of Na+, these results identify mechanistic components underlying ion-substrate coupling in NSS transporters.

## Significance Statement

Membrane transport proteins are responsible for moving substrates such as nutrients, vitamins, drugs and signaling molecules across cellular membranes. A subset of these proteins, the ion-coupled transporters, use a transmembrane ion gradient to drive energetically unfavorable substrate movement from a lower concentration on one side of the membrane to a higher concentration on the other side. They do this by coupling the movement of substrate and ions in the same or the opposite direction. Coupled transport requires conformational changes that occur exclusively or predominantly when specific conditions of ion and substrate binding are met. This work identifies, for a family of Na+-coupled neurotransmitter transporters, how the rules controlling these conformational changes are encoded in the transporter structure.

## Introduction

Ion-coupled transporters have the remarkable ability to transport substrates against their concentration gradient, an energetically uphill process, by utilizing the energy stored in a transmembrane ion gradient. The downhill flux of ions drives substrate transport because transporters catalyze the stoichiometric coupled movement of solutes, either in the same (symport) or opposite direction (antiport). As we develop insight into the conformational changes that transporters undergo to move their substrates across membranes, it becomes essential to understand how these conformational changes are regulated to couple substrate and ion movements. Here, we examine substrate binding in the bacterial amino acid transporter LeuT and its role in coupling substrate-Na^+^ symport.

LeuT is a homologue of neurotransmitter transporters in the NSS (Neurotransmitter:Sodium Symporter, SLC6) family (1). Neurotransmitter transporters in this family are responsible for the re-uptake, after release, of transmitters such as GABA, glycine, norepinephrine, dopamine and serotonin (2, 3). They are targets for antidepressants such as Prozac and Lexapro and drugs of abuse such as cocaine and amphetamines (2). Transporters in this family use the energy from transmembrane gradients of Na^+^ Cl^−^, and K+ and the membrane potential to drive substrate accumulation within cells (4). Almost all NSS transporters are coupled to the Na^+^ gradient, many also to Cl^−^ and at least one to K+. LeuT couples transport of an amino acid substrate to the electrochemical potential of Na^+^ (1, 5). Several other transporter families are structurally similar to LeuT (6), and many of these families also contain members that utilize the electrochemical potential of Na^+^ as a driving force (7). Among these are the SSS (Sugar: Sodium Symporter, SLC7) family containing SGLT Na^+^-sugar symporters (8) and the NCS1 (Nucleobase:Cation Symporter-1) family containing Mhp1 (9).

Transporters like LeuT are thought to work by an alternating access mechanism in which a central binding site is alternately exposed to opposite sides of the membrane by conformational changes that open and close access to that site (Fig. 1A) (10-12). X-ray structures of LeuT provide insight into the conformational changes that occur during transport. They show LeuT in inward- and outward-open conformations (13), and in an outward-occluded conformation with two Na^+^ ions bound (Na1 and Na2) and a molecule of leucine in the substrate site (1, 14, 15) (Fig. S1). Structures of the transporters for dopamine and serotonin (DAT and SERT) reveal these proteins in outward-open conformations quite similar to that of LeuT (16, 17).

**Figure 1.**
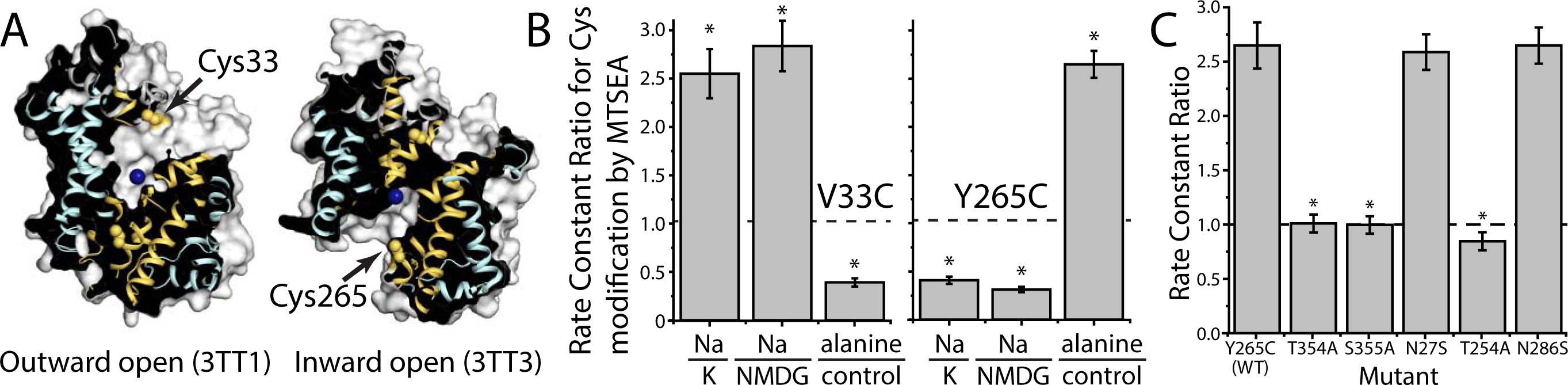
*Na*^+^ *and substrate induce reciprocal changes in LeuT extracellular and cytoplasmic permeation pathways.* A. Location of cysteines used to probe pathway accessibility. Cut-open views of outward-open (PDB 3TT1) and inward-open (PDB 3TT3) LeuT structures (13) are shown from within the membrane plane, with the extracellular side above and the cytoplasm below the structure. The scaffold domain (TMs 3–5 and 8–12) and the bundle domain (TMs 1, 2, 6 and 7) are colored pale cyan and yellow-orange, respectively. The blue sphere in the center of the structures represents the relative position of Na2 in the 3TT1 structure (no Na^+^ was bound in 3TT3). The modified residues V33C and Y265C are shown as spheres, to illustrate the accessibility of V33C to the extracellular pathway, and C265 to the cytoplasmic pathway. B. Accessibility changes induced by Na^+^ and alanine in pathway cysteine mutants. The response of Cys33 in the extracellular pathway (left) and Cys265 in the cytoplasmic pathway (right) to Na^+^ and alanine is shown as a ratio of rate constants for MTSEA modification of the sole cysteine in each mutant (see SI “Modification rates and ratios”). Reactivity of Cys33 was enhanced by 200 mM Na^+^ relative to a control in either Κ+ or NMDG+ and was decreased by addition of 10 μΜ alanine in 10 mM Na^+^ relative to control, Na^+^ alone (left column). The reverse pattern of responses was observed in Y265C (right), consistent with the expectation that only one pathway can be open at any one time. Individual rates are shown in Figure S2A. The dashed line in this and other figures represents a ratio of 1.0 indicating no change in response to alanine (or Na^+^). Asterisks indicate ratios significantly different (P < 0.05) from 1 using Student’s T-test. All error bars represent the Standard Error of the Mean (SEM); n ≥ 3. C. Accessibility changes induced by alanine in Na1- and Na2-site mutants. Rate constant ratios for alanine addition relative to a control in the presence of Na^+^ alone. Alanine was added at concentrations determined to be saturating for each mutant. Individual rates are shown in Figure S2B. MTSEA concentrations were 0.1 μΜ - 0.1 mM for N286S, 0.05 μΜ - 0.5 mM for S355A and 0.1 μΜ - 1 mM for all other mutants. Asterisks indicate values significantly different (P < 0.05) from that of Y265C.

In the alternating access model for ion-coupled transport (10-12) the conformational changes that move substrates and ions across the membrane must be regulated by binding of ions and substrates. For a Na^+^-amino acid symporter like LeuT, these conformational changes should be allowed when both Na^+^ and amino acid substrate are bound or when both binding sites are empty, but not when only Na^+^ or only substrate are bound (18). Together, these rules prevent uncoupled movement of Na^+^ or substrate.

Observations with the homologous bacterial tyrosine transporter Tyt1 hint at the coupling rules for this family. Specifically, Quick et al. (19) measured the accessibility of cysteines in the pathway between the central binding site of Tyt1 and the cytoplasm. The cysteines became less accessible in the presence of Na^+^, suggesting that Na^+^ closed this cytoplasmic pathway. Tyrosine, which had no effect in the absence of Na^+^, was able to reverse the effect of Na^+^ on cysteine accessibility. These results were interpreted to mean that Na^+^ stabilized the transporter in an outward-open conformation in which the cysteines were buried, and that tyrosine binding overcame the effect of Na^+^, allowing a transition to the inward-open form in which the cysteines were exposed and from which Na^+^ and substrate dissociate to the cytoplasm.

Similar effects of Na^+^ and substrate have been observed in other NSS transporters (20-27), suggesting that they reflect the coupling rules in all members of this family. The general features of this mechanism are that, first, Na^+^ binding stabilizes the transporter in an outward-open conformation, preventing uncoupled Na^+^ transport. Second, an interaction between the amino acid substrate carboxyl group and the Na1 site Na^+^ ion renders substrate binding strongly dependent on Na^+^; this minimizes the possibility that substrate could be transported without Na^+^ (25, 28). And finally, when Na^+^ is bound, the binding of substrate allows transition from an outward-open, through an occluded (29), to an inward-open conformation, leading to release of Na^+^ and substrate intracellularly.

Shi et al. (28) proposed that occupation of a second site in the extracellular pathway, named S2 (Fig. S1), is required for the ability of substrate to convert LeuT to an inward-open state. Evidence favoring this proposal has come mainly from binding and displacement studies and from steered molecular dynamics (MD) simulations (15, 28, 30). Mutation of residues proposed to contribute to the S2 site, Ile111 and Leu400, inactivated transport, highlighting the importance of this region (28). In addition, a Leu400 mutant was unable to undergo substrate-induced conformational changes (22). However, crystallographic studies have not detected substrate binding in that site (31-33) and more recent studies using MD simulation (34) and solid-state NMR (35) have argued against substrate binding in the S2 site.

In spite of these significant efforts, it remains unclear which structural elements in LeuT are responsible for encoding the coupling rules in LeuT and ensuring its responses to substrate binding. To identify these elements, we tested the effect of mutating residues at both Na^+^ and substrate binding sites. Specifically, we determined the effect of Na^+^ and substrate on the conformation of LeuT mutants in *E. coli* membranes by measuring the reactivity of cysteine residues in the substrate permeation pathways. We characterized substrate binding in these mutants using the scintillation proximity assay with solubilized LeuT (36) and determined transport using purified LeuT reconstituted in proteoliposomes. In previous work (25), we identified Na2 as the site at which Na^+^ binding stabilizes the outward-open conformation. This site is at the interface of the scaffold and bundle domains (Fig. S1), which move relative to one another during the transition between inward- and outward-open conformations (13). Here, we identify a network of interactions that connects the two domains when substrate binds at the central binding site. Importantly, we also identified specific interactions central to the substrate’s ability to overcome the effect of Na^+^, close the extracellular pathway, and open the cytoplasmic pathway.

## Results

*Alanine reverses the conformational effect of sodium.* To measure conformational changes in LeuT, we utilized an assay based on accessibility of a cysteine residue, strategically placed in one of the two substrate permeation pathways (Fig. 1A). This cysteine reacts with the aqueous cysteine reagent MTSEA (2-aminoethyl methanethiosulfonate hydrobromide) faster when that pathway is open (25, 37). The positions were selected based on previous studies demonstrating conformation-dependent reactivity of cysteine residues in the cytoplasmic (19, 38) and extracellular (39) permeation pathways of other NSS transporters. Figure 1B shows that this assay is able to detect conformational changes in response to Na^+^ and substrate (alanine, a good substrate for LeuT (14)). The conformational response in this figure is shown as the ratio of rate constants in the presence and absence of Na^+^ or alanine. This ratio would be 1 (dashed line) if a treatment had no effect on LeuT conformation and thus no effect on cysteine reactivity. Na^+^ increased the accessibility of Cys33 in the extracellular pathway as indicated by a ratio >1 (Fig. 1B, left). On the other hand, Na^+^ decreased the accessibility of Cys265 in the cytoplasmic pathway, leading to a ratio <1 (Fig. 1B, right). Alanine reversed this effect of Na^+^, decreasing accessibility in the extracellular pathway (Fig. 1B, left), and increasing accessibility in the cytoplasmic pathway accessibility (Fig. 1B, right), as indicated by the rate constant ratios. Data from these two mutants, V33C and Y265C, suggest that the conformational changes induced by Na^+^ and substrate result in inverse accessibility changes for the two pathways. Individual rate constants are shown in Figure S2A. A cysteine inserted at a control position on the extracellular surface, H480C (40), showed no change in reactivity in response to Na^+^ relative to NMDG (ratio = 0.98 ± 0.10) or in response to alanine in the presence of Na^+^ (ratio = 1.02 ± 0.35).

These data show that substrate, in the presence of Na^+^, increases the population of transporters with the cytoplasmic pathway open, as observed previously for LeuT and the homologous Tyt1 (19, 22), and that this opening is associated with a decrease in the population of protein with the extracellular pathway open. This phenomenon illustrates a corollary of the alternating access hypothesis (10), namely that both pathways should not be open simultaneously to avoid forming a channel that would dissipate transmembrane ion and substrate gradients. Furthermore, the ability of Na^+^ to close the cytoplasmic pathway and open the extracellular pathway was evident whether K+ or NMDG+ (N-methyl-D-glucamine) was used as the control cation (Fig. 1B), contrary to a recent suggestion that K+, rather than Na^+^, was responsible for conformational change (41). We do not understand the reason for this discrepancy. However, the previous study used a LeuT mutant in which two residues were mutated to histidine to coordinate a Ni^2^+ ion, and another residue mutated to cysteine to attach a fluorophore. These mutations were made so that conformational changes could be measured, in detergent, by transition metal FRET (41). In contrast, our constructs contained only one mutation, to cysteine, and reactivity was measured in *E. coli* membranes in the absence of detergent. We believe, therefore, that our measurements were less prone to artifacts.

*The conformational responses of LeuT are not due to a transmembrane concentration gradient.* Na^+^ and substrate influence the conformation of LeuT in the absence of transmembrane concentration gradients of these ligands, as evidenced by previous studies using detergent-solubilized LeuT (20-22). However, the possibility that being in a sealed vesicular structure would influence the conformational response of LeuT has not previously been tested. Our experiments used detergent-free *E. coli* membranes prepared using a cell disruptor to assay conformational change. We observed these membranes to be severely deficient in their ability to generate and maintain concentration gradients, as if they were leaky, non-vesicular, or sealed but topologically inverted (Fig. S3). Indeed, these membranes were unable to accumulate [^3^H]proline in the presence of D-lactate (42), unlike vesicles from the same cells prepared by osmotic shock. To test the effect that a substrate gradient would have on accessibility measurements, we used the detergent lauryl-maltose-neopentyl-glycol (LMNG) in an amount just sufficient to permeabilize the transport-competent osmotic shock vesicles expressing LeuT Y265C (*inset* A) and compared the ability of alanine to induce a LeuT conformational change in LMNG-permeabilized and intact vesicles. As shown in *inset* B, there was no difference in the response to alanine, indicating that the conformational response to substrate was not due to a transmembrane substrate gradient.

*The response of LeuT to alanine requires residues in the sodium binding sites.* We previously demonstrated that mutation of the Na2, but not the Na1, site of LeuT blocked the conformational response to Na^+^ (25). Here, we examined the effect of amino acid substrate when the Na1 or Na2 sites were mutated. In the background of Y265C, Na2 site mutations T354A and S355A, which were previously shown to eliminate the response to Na^+^, also blocked the alanine-induced conformational change observed in the presence of Na^+^ (Fig. 1C, ratios, and S2B, rates). This is consistent with the idea that Na^+^ does not bias the transporter toward outward-facing conformations when Na2 is disrupted (25), and, consequently, there is no reversal by alanine. Na1 site mutants N27S and N286S responded to Na^+^ (25) and also responded to alanine in the presence of Na^+^ (Fig. 1C). On the other hand, T254A, although shifted by Na^+^ toward outward-facing conformations (25), was not affected by saturating concentrations of alanine. T254A had low affinity for leucine (25), but that alone does not account for the lack of substrate response because the alanine response of N27S was normal despite having an even lower leucine affinity than T254A (25). T254 is uniquely positioned to mediate the interaction between Na1 and E290, which was proposed to be involved in closing the extracellular pathway (43). Thus, it is likely that the T254A mutation blocks the response to substrate because it disrupts this interaction of Na1 with E290. Overall, these results indicate that the ability to respond to Na^+^ is apparently required for substrate to induce conformational change.

*Mutation of the central substrate site identifies a critical interaction.* We reasoned that substrate binding must facilitate interactions at its binding site (Fig. 2A) to overcome the conformational restriction imposed by Na^+^. To identify those interactions, we mutated residues that interact with amino and carboxyl groups of the amino acid substrate, as well as with the side chain of the substrate (Fig. 2A, Fig. S1 shows the position of mutated sites in the LeuT structure). In molecular dynamics simulations of LeuT structures in the outward-occluded state (Fig. S4A), the substrate carboxyl group coordinates Na1 and is hydrogen-bonded by the hydroxyl group of Tyr108 and the main chain amide nitrogen of Leu25 (Fig. 2A). The substrate amino group hydrogen bonds to Ser256 and to main chain oxygens from Phe253 and Thr254, while the side chain pocket is lined with several hydrophobic and aromatic side chains including those of Tyr108 and Ile359 (1, 14) (Fig S4A). Notably, however, the effect of modifying these central binding site residues on substrate affinity had not previously been measured, except for Y108F (31). Accordingly, we mutated Tyr108 to Phe, Ser256 to Ala and Ile359 to Ala, all in the Y265C background, to disrupt interactions between LeuT and the substrate amino and carboxyl groups and its side chain. We used the scintillation proximity assay (SPA) (36) to measure the binding affinity for [^3^H]leucine to LeuT solubilized from *E. coli* membranes and bound to Copper YSI beads through a C-terminal Hiss tag. We also used this assay to determine affinity for unlabeled alanine by displacement of [^3^H]leucine, using a Na^+^ concentration that was half-maximal for leucine binding. Each of the binding curves was best fit with a single affinity and all of the substrate site mutations greatly increased the K_D_ for leucine and alanine binding (Table 1 and Fig. 2B). In none of these mutants did we detect measurable binding with the high affinity characteristic of WT or Y265C. From these measurements, we were able to identify alanine concentrations sufficient to saturate the binding site for each mutant, namely 0.2 mM alanine for I359A, and 5 mM for Y108F and S256A.

**Figure 2.**
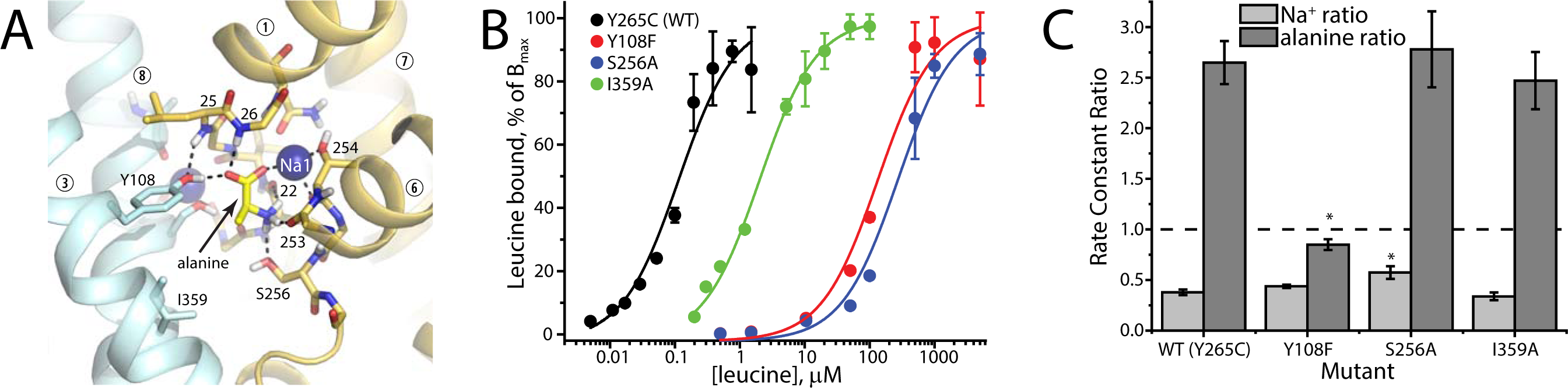
*Substrate binding site mutations identify a role for the substrate carboxyl and Tyr108 in conformational change.* A. Close-up of the central binding site in simulations of outward-occluded LeuT. The substrate carboxyl group coordinates Na1, and accepts hydrogen bonds from Tyr108 from TM3 in the scaffold domain and the backbone amide of Leu25 in the bundle domain. The substrate amino group donates hydrogen bonds to Ser256 and main chain carbonyl oxygen atoms from Phe253 and Thr254, from the bundle domain (coloring as in Fig. 1). The substrate side chain lies in a hydrophobic pocket formed by several residues, including Ile359 from TM8 in the scaffold. Only polar hydrogen atoms are shown explicitly, for clarity. B. [^3^H]Leucine binding to detergent-solubilized LeuT. Leucine binding was measured using the SPA assay to determine the effect of mutation on substrate affinity. Compared with the control (Y256C), mutation of Ile359 increased leucine Kd 12-fold, Y108F increased Kd 1000-fold and S256A increased Kd 2000-fold (see Table 1). Similar changes were found for the ability of alanine to displace [^3^H]leucine. In none of these mutants did we observe a binding component that would correspond to a high-affinity binding site at S2; n=3. C. Accessibility changes induced by Na^+^ and alanine in central binding site mutants show that Tyr108 is critical for substrate-but not Na^+^-dependent conformational change. The light gray bars show that mutation of Y108, S256 and I359 had only minor effects on the ability of 200 mM Na^+^ to decrease Cys265 accessibility relative to controls with equimolar NMDG+ (or Κ+ in some experiments). The dark gray bars show the effect of the alanine addition in presence of Na^+^, relative to controls with Na^+^ alone. Alanine increased C265 accessibility in mutants S265A and I359A, similar to the Y265C control, but Y108F was unable to respond to alanine. We used 100 mM Na^+^ and 5 mM alanine for S265A and Y108F, 50 mM Na^+^ and 200 μΜ for I359A. Asterisks indicate values significantly different (P < 0.05) from those of Y265C; n=4 except S256A ± alanine, n=3.

**Table 1.** Summary of binding and transport measurements for LeuT mutants. The K_05_ value for Na^+^ represents the concentration of Na^+^ that supported half-maximal binding at 1 μM [^3^H]leucine. Subsequent measurements of leucine binding and displacement by alanine were performed using the K_05_ concentration of Na^+^ for each mutant. All constructs contained a His_8_ tag at the C-terminus. Except for wild type and V33C, all constructs contained the Y265C mutation. ND: Not detected. For binding measurements, n=3; for transport, n=2.

| Position | $K_{0.5}\text{-Na}$, mM | $K_D\text{-leu}$, $\mu\text{M}$<br>@ $K_{0.5}\text{-Na}$ | $K_I\text{-ala}$, $\mu\text{M}$<br>@ $K_{0.5}\text{-Na}$ | Influx,<br>$\text{pmol mg}^{-1}\text{ min}^{-1}$ |
| --- | --- | --- | --- | --- |
| **Wild Type** | $11.2 \pm 1.4$ | $0.075 \pm 0.004$ | $2.56 \pm 0.51$ | $327 \pm 80$ |
| **Y265C** | $17.0 \pm 2.3$ | $0.176 \pm 0.079$ | $2.04 \pm 0.32$ | $339 \pm 66$ |
| **V33C** | $3.06 \pm 1.78$ | $0.075 \pm 0.013$ | $11.1 \pm 4.7$ | $448 \pm 52$ |
| **Central binding site mutants** |  |  |  |  |
| **Y108F** | $71.2 \pm 5.4$ | $105 \pm 8$ | $302 \pm 29$ | ND |
| **S256A** | $113 \pm 9$ | $231 \pm 8$ | $844 \pm 32$ | ND |
| **I359A** | $51.7 \pm 7.8$ | $1.24 \pm 0.03$ | $21.5 \pm 1.9$ | $22 \pm 1$ |
| **Extracellular pathway mutants** |  |  |  |  |
| **L29M** | $21.4 \pm 5.6$ | $0.260 \pm 0.075$ | $8.70 \pm 1.98$ | $371 \pm 74$ |
| **I111L** | $18.4 \pm 4.9$ | $0.142 \pm 0.036$ | $4.67 \pm 1.10$ | $15 \pm 9$ |
| **A319M** | $2.03 \pm 0.4$ | $0.100 \pm 0.026$ | $9.50 \pm 2.16$ | ND |
| **L400A** | $2.03 \pm 0.4$ | $0.0784 \pm 0.020$ | $3.78 \pm 0.91$ | ND |
| **D401V** | $4.30 \pm 0.9$ | $0.149 \pm 0.040$ | $6.77 \pm 1.56$ | ND |

The above results demonstrate the crucial role of these central site residues in substrate binding, even when Na^+^ is present. However, it remains unclear whether these residues are necessary only for high-affinity binding, or also for the conformational changes required for transport. To determine the effect that these mutations would have on conformational change, we tested, for each mutant, the ability of Na^+^ to decrease Cys265 accessibility and of alanine to reverse the effect of Na^+^. The results, shown as ratios of reaction rates ± Na^+^ or ± alanine in the presence of Na^+^ (Fig. 2C; individual rate constants in Figures S5A and S5B), demonstrate that each of the mutants responds to Na^+^ by closing the cytoplasmic pathway, similar to Y265C. In both S256A and I359A, this pathway opens in response to alanine, as happens with Y265C. For Y108F, however, alanine has no significant effect on conformation, even though this mutant responds normally to Na^+^. These results identify the hydroxyl group in Tyr108 as a critical moiety for interactions in LeuT that allow binding of amino acid substrate to induce conformational change.

*The extracellular pathway participates in substrate dependent conformational change.* Singh et al. (44) identified a ligand binding site in the extracellular pathway and Shi et al. (28) proposed that a second molecule of substrate, binding at this S2 site (Fig. 3A), was required to allosterically trigger Na^+^ and substrate release to the cytoplasm. Mutating either of two residues (Ile111 and Leu400) in this S2 region was shown to inactivate alanine transport (28) and mutation of Leu400 was shown to block the effect of alanine on conformational dynamics (22). To evaluate the role of S2 on Na^+^ and substrate-induced conformational change, we mutated several residues in this region (Fig. 3A and S1) and studied the effect of mutation on the conformational response of LeuT-Y265C. In addition to Ile111 and Leu400, we mutated two residues, Leu29 and Ala319, whose side chains are close to non-protein electron densities in the LeuT inward-open structure (13), as well as Asp401, which is outside the proposed S2 binding site, but is hydrogen-bonded by the main chain amide of Ala319 across the pathway (Fig. 3A, S4B).

**Figure 3.**
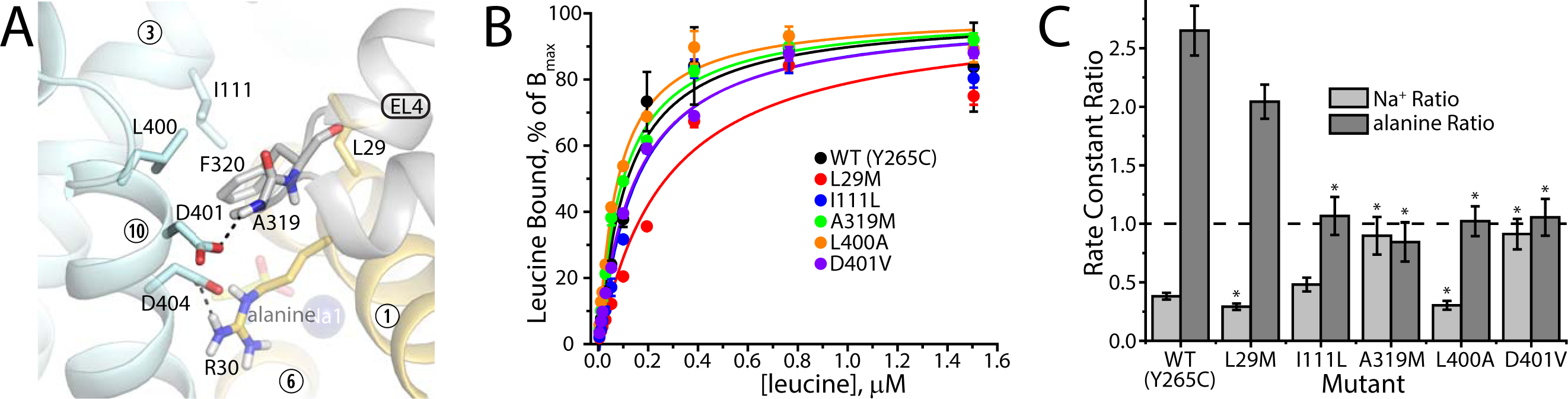
*Extracellular pathway residues contribute to substrate dependent conformational change.* A. Close-up of the LeuT extracellular pathway in simulations of the inward-open conformation (PDB 3TT3), in which Ile111, Ala319, Phe320 and Leu400 form a network of Van der Waals contacts. Dashed black lines indicate an ionic interaction between Asp404 and Arg30 and a hydrogen bond between Asp401 and Ala319 from extracellular loop 4 (EL4). Aside from Ala319 and Phe320, backbone atoms are not shown for clarity. B. Extracellular pathway mutations have small effects on leucine affinity. [^3^H]Leucine binding was measured using a Na^+^ concentration giving half-maximal binding at 1 μΜ leucine for each mutant (see Table 1); n=3. C. Accessibility changes induced by Na^+^ and alanine in extracellular pathway mutants suggest that scaffold-bundle interactions in the extracellular pathway are required for Na^+^ and substrate-dependent conformational change. Of the extracellular pathway mutants, only L29M, which was fully functional for alanine transport (Table 1) responded to both Na^+^ and alanine like the background construct (Y265C). All the other mutants were blocked in their response to either alanine (I111L and L400A) or both Na^+^ and alanine (A319M and D401V). All mutations were in the Y265C background. To measure the response to alanine, we used 20 mM Na^+^ for L29M and I111L, 2 mM Na^+^ for A319M and L400A, and 5 mM Na^+^ for D401A. Alanine concentrations were 50 μΜ for L29M and A319M, 30 μΜ for I111L and L400A and 70 μΜ for D401A. Asterisks indicate values significantly different (P < 0.05) from those of Y265C. For L29M, I111L and D401V ± Na^+^, n=5; for WT (Y265C), L400A and A319M, n=4; for D401V ± alanine, n=3.

In contrast to mutations at the central substrate site, mutations in the S2 region did not dramatically influence [^3^H]leucine binding affinity, its dependence on Na^+^ concentration, or its displacement by alanine (Fig. 3B, Table 1). The largest affinity decrease was observed for L29M and A319M, which decreased alanine affinity less than 5-fold relative to Y265C alone (Table 1). Leucine binding to several of the mutants, A319M, L400A and D401V, was more sensitive to Na^+^, and all of the mutants bound alanine with moderately decreased affinity (2- to 5-fold), but leucine affinity was hardly affected by mutation (Fig. 3C and Table 1). By comparison, mutations at the central substrate site decreased leucine and alanine affinity by 12- to 2000-fold (Fig. 2B and Table 1).

Although the above data suggests that the S2 region is not directly involved in substrate binding, the deleterious effect of I111C and L400S on transport (28) and conformational dynamics (22) suggests that closure of the extracellular pathway may be sensitive to its composition. However, the previous studies did not indicate whether these mutants were capable of conformational change. We therefore tested the ability of Na^+^ and alanine to alter the cytoplasmic pathway accessibility in S2 mutants. Figure 3C clearly shows that mutation of Ile111 and Leu400 does not prevent their conformational response to Na^+^, which is similar to that of Y265C. However, in the presence of Na^+^, I111L and L400A were unresponsive to saturating concentrations of alanine (i.e., ± alanine ratio = 1), indicating that these residues participate in interactions required specifically for substrate to overcome the conformational effects of Na^+^ binding. Of the three remaining mutants, A319M and D401V responded to neither Na^+^ nor alanine. In the case of A319M, the addition of the 2-thiapropyl group to the alanine side chain apparently prevented of the extracellular pathway from closing. The measured rates of Cys265 modification in A319M were far lower than any other LeuT mutant we have studied (Figs. S6A and S6B). We surmise that the additional bulk added to the Ala319 side chain cannot be accommodated when the extracellular pathway closes, strongly destabilizing inward-open conformations. Asp401 is hydrogen bonded by the amide nitrogen of Ala319 in simulations of the inward-open conformation (Fig. 3A, S4B). While Asp401 was not considered part of the S2 site (28), it provides interactions between bundle and scaffold domains in the extracellular pathway. As mentioned, this mutant responded to neither Na^+^ nor alanine (Fig. 3C), although accessibility measurements showed that the cytoplasmic pathway was more accessible than in A319M (Figs. S6A and S6B). Only L29M showed a conformational response similar to that of Y265C for both Na^+^ and alanine (Fig. 3C), consistent with its position on the edge of the network of interactions (Fig. 3A).

*Effect of substrate site and S2 mutations on transport activity.* Reconstitution of LeuT mutants in proteoliposomes allowed us to test their ability to catalyze alanine transport. The single-Cys mutants Y265C and V33C, and the L29M mutant in the Y265C background all had activity close to that of WT LeuT (Table 1). All of the mutants in which either the Na^+^-dependent or substrate-dependent conformational changes were inhibited (Y108F, I111L, A319M, L400A and D401V) were severely defective in transport, although I111L had a trace of residual transport activity (Table 1). The central site mutant I359A retained a low level of transport activity. The relatively low activity of this mutant was likely due to poor substrate affinity, which was verified by the observation that 2 and 10 μM unlabeled alanine decreased I359A transport of [^3^H]alanine by 3 and 16% respectively, compared to 60 and 80% for LeuT-Y265C. These results are consistent with a K_m_ increase of >6-fold for I359A. Structures with leucine bound (e.g. PDB 2A65) suggest a direct interaction of the substrate with I359, consistent with the effect on [^3^H]leucine binding (Table 1). However, when the substrate is the smaller alanine, its side chain methyl group is ~4-6 Å from I359, which is longer than a direct van der Waals interaction (Fig. 2A and S4A). We attribute the strength of the effect on alanine transport to an additional cavity created in I359A, which we expect would be more hydrated, thereby disfavoring alanine binding. However, we cannot rule out that the I359A mutation also influences binding to the inward-facing conformation, for which a substrate-bound structure is not yet available.

We were unable to measure transport activity using proteoliposomes containing the S256A mutant, despite its normal conformational responses to Na^+^ and alanine (Fig. 2C). We considered the possibility that the return step in transport, when apo-LeuT converts from inward- to outward-open, was defective in S256A. To bypass the apo return step, we carried out a counterflow experiment (Fig. S7) in which LeuT proteoliposomes were loaded with unlabeled alanine. This measurement showed an accumulation of intravesicular [^3^H]alanine by WT but not by S256A (Fig. S7), suggesting that the mutant did not catalyze even this partial reaction. However, the affinity of S256A for alanine was 500-fold lower than that of Y265C, which might cause the transport defect (Table 1). Indeed, the I359A mutation, which decreased alanine affinity only 8.4-fold, decreased transport activity 15-fold. We therefore consider it likely that the decreased substrate affinity of S256A was responsible for our inability to detect transport. However, we cannot rule out the possibility that S256A, or some other mutants might be structurally more labile during the reconstitution procedure, which would also lead to activity loss.

*Conformational changes in SERT.* The mammalian transporter SERT also belongs to the NSS family, but has several significant differences from LeuT. Most important for substrate-induced conformational changes is that its substrate, serotonin (5-HT), lacks the carboxyl group present in LeuT substrates. In this way, SERT is similar to the catecholamine transporters NET and DAT, which transport norepinephrine and dopamine, respectively. However, these three amine transporters all contain an aspartate residue in TM1 (Asp98 in SERT) at the position of a highly conserved glycine (Gly24 in LeuT) in NSS amino acid transporters (Fig. 4A).

**Figure 4.**
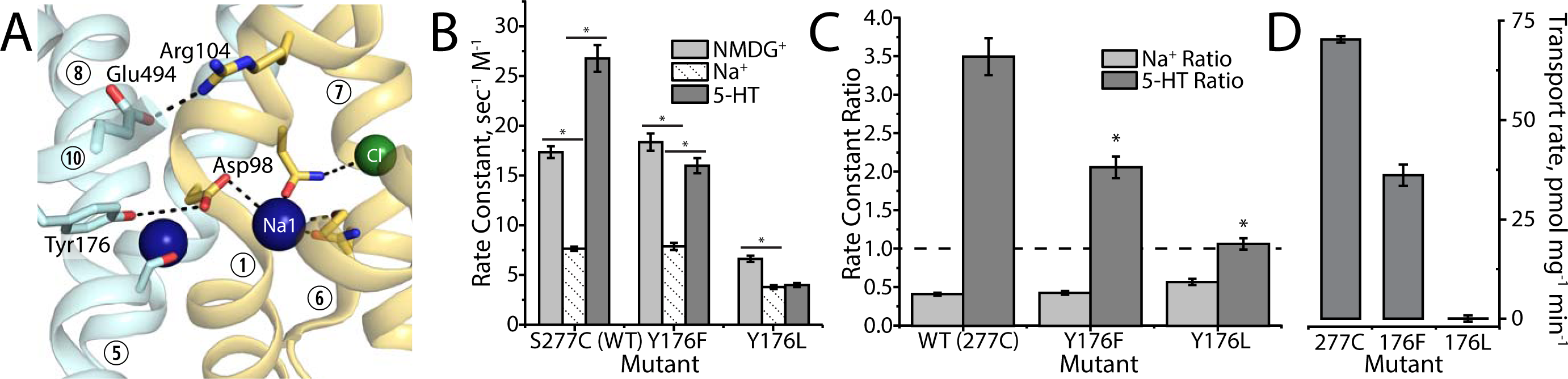
*Disruption of an ionic network in SERT blocks substrate-dependent conformational change.* A. The central binding site in thermostabilized human SERT bound to citalopram (PDB 5I71), shown after removal of citalopram. Binding site residues are in stick representation, and Na^+^ and Cl^−^ ions are shown as spheres. The β-carboxyl group of Asp98 in SERT is in a position corresponding to the substrate carboxyl in LeuT and connects Tyr176 (which corresponds to LeuT Tyr108) with Na1. B. Individual rates of SERT Cys277 modification by MTSEA in Tyr176 mutants in 150 mM NMDG-Cl (light gray), equimolar NaCl (diagonal pattern) and NaCl + 10 μΜ 5-HT (dark gray). The modification rate for Y176L in the absence of Na^+^ and 5-HT was decreased relative to the S277C control, suggesting that this mutant was somewhat biased toward inward-closed states. For Y176F and Y176L, n=3; for S277C, n=4. Asterisks indicate rate constants significantly different (P < 0.05) from those in NaCl. C. 5-HT-dependent conformational change in SERT is inhibited by mutation of Y176. Modification of the cytoplasmic pathway residue Cys277 by MTSEA was decreased by 150 mM NaCl, relative to NMDG-Cl (light gray columns), and enhanced by 10 μΜ 5-HT in the presence of 150 mM NaCl, relative to NaCl alone (dark gray columns), as shown previously (24, 27, 38). These changes indicate a decrease and increase, respectively, in cytoplasmic pathway accessibility (38). When Y176 was mutated to leucine, Na^+^ still decreased accessibility, but 1 mM 5-HT had no effect on opening the cytoplasmic pathway (ratio = 1). SERT Y176F responded to Na^+^ like the control, but had a decreased response to 10 μΜ 5-HT. Asterisks indicate values significantly different (P < 0.05) from those of S277C control. D. 5-HT transport rates in Tyr176 mutants (n=3). Y176F was approximately half as active for transport as the control, and Y176L was inactive for transport. 5-HT affinity was similar in Y176F and the background construct (S277C) but was decreased in Y176L and D98N; Ki values for 5-HT displacement of β-CIT were 1.13 ± 0.05 μΜ for S277C, 1.18 ± 0.05 μΜ for Y176F, 65 ± 4 μΜ for Y176L and 500 ± 9 μΜ for D98N (n=2).

Because the presence of Asp98 would allow SERT to form interactions analogous to those in the network critical for substrate-dependent conformational change in LeuT (Tyr108…substrate-COO^−^…Na1), we mutated the corresponding tyrosine (SERT Tyr176) and Asp98 in rat SERT. Both these residues were previously shown to be critical for transport by SERT (45, 46), but the affected step in transport had not been determined. Using the accessibility of a cysteine residue (Cys277) in the cytoplasmic permeation pathway (24, 38), we measured the effect of mutating these two residues on conformational changes induced by 5-HT and Na^+^.

In SERT, the D98N mutant was inactive for transport and had much lower affinity for 5-HT and high-affinity SERT ligands: β-CIT binding was undetectable and citalopram binding was < 7% of control. Consequently, we were unable to measure Cys277 accessibility changes in response to Na^+^ or 5-HT in D98N. Mutation of Tyr176 to Phe decreased transport about 50%, and Y176L was completely inactive, like Y108F in LeuT (Fig. 4D), but both mutants retained ligand binding activity. Both Y176F and Y176L responded normally to Na^+^, which decreased accessibility of Cys277 as it did in the background construct, indicating no defect in the ability of Na^+^ to stabilize the outward-open conformation of SERT (Fig. 4C, rates in Fig. 4B). A Y156F mutant in DAT was predicted by MD simulation to prefer more outward-open conformations (47) although we saw no such effect in the corresponding Y176F SERT mutant (Fig. 4B). The ability of 5-HT to convert SERT to an inward-open state (measured by the increased Cys277 accessibility) was also lower in Y176F and undetectable in Y176L (Fig. 4B, C), corresponding with the decreased transport activity of these mutants (Fig. 4D). These results indicate the importance of Tyr176 for substrate-induced conformational change in SERT.

## Discussion

This work identifies structural elements in LeuT that are required for the conformational response to substrate binding. In effect, these elements, together with Na^+^ interactions at the Na2 site (25), facilitate control of LeuT conformation by ion and substrate binding events. These binding site interactions, therefore, encode the rules essential for ion-substrate coupling in LeuT. Specifically, our results point to the critical importance, in LeuT, of a network of interactions involving Y108, Na1, and backbone atoms, mediated by the substrate carboxyl group (Fig. 2A). This network forms upon substrate binding, but is not required for binding. Instead, its role appears to be to stabilize the close proximity of the extracellular halves of the scaffold and bundle domains of LeuT, which become progressively closer in structures moving from outward-open to outward-occluded to inward-open conformations. We propose that the substrate acts as a bridge between Tyr108 in the scaffold and Na1 in the bundle, and that Tyr108 contributes to stabilize a configuration where substrate “holds” the two sides of the extracellular pathway together, acting together with residues in the pathway (Figs. 3, S1). In this way, Y108, the substrate carboxyl group and Na1 mediate the conformational change required to close the extracellular pathway, thus allowing the cytoplasmic permeation pathway to open. As mentioned, this substrate-dependent effect is one of two conformational changes identified by Quick et al (19) in the homologous Tyt1, the other being a shift toward the outward-open conformation imposed by Na^+^ binding at the Na2 site (20-23, 25). Together, these two conformational changes define the rules that enforce the stoichiometric coupling of Na^+^ and substrate flux in LeuT and other NSS family transporters.

We show here that Tyr108 is required for the conformational change in LeuT that allows substrate and Na^+^ release to the cytoplasm. This same conformational change in SERT requires the corresponding residue Y176, strongly implying that a network of interactions linking TM3 and Na1 is essential for transport throughout the NSS family. In agreement with this interpretation, NSS amine transporters, whose substrates lack the critical carboxyl group of NSS amino acid substrates, contain an aspartate in TM1 (D98 in SERT), which is unique to the NSS transporters whose substrates lack carboxyl groups. The carboxyl group of this aspartate is located, in outward-open X-ray structures of SERT and DAT, between the TM3 tyrosine and Na1 (Fig. 4A), like the substrate carboxyl in LeuT (1, 17, 48). Nevertheless, the NSS amine transporters must use a variation of the LeuT mechanism, since substrate is still required for conformational change (24) despite the presence of a carboxyl group supplied by the protein. The residual activity of SERT Y176F was not seen in LeuT with the corresponding Y108F mutation, suggesting the importance of side chain interactions between 5-HT and Tyr176, perhaps involving cation-π or π-electron stacking of the rings. In the context of SERT, but not LeuT, an electropositive phenolic hydrogen atom of Phe176 may partially replace the WT tyrosine hydroxyl group in Y176F. Furthermore, Tyr176 likely interacts with the fixed γ-carboxyl of Asp98 in SERT, rather than with the carboxyl group of a dissociable substrate molecule as in LeuT. This difference may also help explain why substitution with Phe is less deleterious in the SERT context. Future studies will address the mechanism used in NSS amine transporters to activate the interaction between TM3 and Na1 when substrate binds.

An earlier study of LeuT proposed that substrate binding at a site (S2) in the extracellular pathway, induced conformational changes leading to cytoplasmic release of substrate and ions (28). In agreement with their results, we show that mutations in and near the S2 region disrupt the conformational changes induced by Na^+^ and substrate binding. However, we prefer to interpret our results as evidence for interactions between bundle and scaffold domains in the extracellular pathway rather than substrate binding at S2 for the following reasons:

1) Our binding measurements are consistent with a single high-affinity substrate site. Mutations in the S2 region had only minor effects on leucine and alanine affinity (Fig. 3B), consistent with either a single site or two independent sites with equal affinity as proposed by Shi et al. (28). However, mutating residues in the central substrate binding site (S1) increased the Kd for leucine and Ki for alanine binding between 12- and 2000-fold with no residual (S2) high-affinity binding detected, despite the prediction (30) that binding at S2 is equal in affinity to S1 and should be unaffected by mutation at S1 (Fig. 2B). Our binding measurements were performed at the same DDM concentration used by Shi et al. (28) to avoid displacement by detergent of leucine bound in the S2 region (15).

2) Increasing the length of a sidechain where substrate was proposed to bind in the S2 locus prevented the extracellular pathway from closing. The effect of mutating Ala319 to Met was to make Cys265 in the cytoplasmic pathway almost inaccessible to modification by the aqueous cysteine reagent MTSEA (Figs. S6A, S6B), an indication that the cytoplasmic pathway was essentially never open. We conclude from this result that the extracellular pathway cannot close in A319M because there is no space in the outward-closed conformation of LeuT for the methionine side chain. This mutant was designed so that the M319 side chain would fit into the proposed S2 binding site, although the flexibility of the side chain could allow it, alternatively, to occupy a non-protein electron density observed near Ala319 in the inward-open X-ray structure of LeuT (SI Figure 12a in ref.(13)). Our results strongly suggest that neither of these sites accommodated the Met319 side chain, supporting the conclusion by Krishnamurthy and Gouaux (13) that there is no room in the closed extracellular pathway for an even bulkier second substrate molecule.

Incidentally, the finding that disrupting interactions in the extracellular pathway prevents opening of the cytoplasmic pathway fulfills an expectation of the alternating access model. In this model, opening one pathway requires that the other pathway be closed to ensure that both pathways are not open simultaneously. The loss of Cys265 reactivity, therefore, argues against the recent proposal (40) that apo-LeuT can adopt a conformation with both pathways open.

In addition to steric disruption by the side chain of Met319, this mutant may also have a diminished ability to undergo a change in backbone dihedral angle, which was observed in comparisons of the outward and inward-facing states in molecular dynamics simulations of LeuT (49). Mutations outside of the S2 site had a similar effect as S2 mutations on substrate-dependent conformational change. Specifically, D401, in the scaffold domain outside the proposed S2 binding site, H-bonds with backbone atoms of EL4, and this interaction appears to be partly responsible for closing the extracellular pathway. Mutation of D401 disrupted conformational changes induced by Na^+^ and substrate (Fig. 3C). Overall, our conclusions are consistent with other recent results that argue against substrate binding at the S2 site (34, 35).

If substrate does not bind in S2, why is substrate-dependent conformational change disrupted by mutation of I111 and L400? This region is clearly important for substrate effects, even if it does not bind a second molecule of substrate. Not only at I111 and L400, but also mutations at A319 and D401 have dramatic effects on substrate-dependent conformational change. We infer that hydrophobic and van der Waals forces (for I111, A319 and L400), H-bonding (for D401) and electrostatic interactions (between R30 and D404 (Fig. S4B), see Ben-Yona and Kanner (50)) are required to stabilize the closed state of the extracellular pathway, although in the presence of sodium, these forces are not sufficient to close the pathway without the action of substrate linking Y108 and Na1 at the central binding site. In agreement with this interpretation, mutation of Thr254 also blocked the effect of substrate on the conformational change (Fig. 1C). This residue was proposed as part of a network of interactions responsible for formation of the R30-D404 salt bridge (43).

In addition to the NSS family, several other transporter families adopt a fold similar to LeuT despite negligible sequence similarity. This larger structural superfamily (51) is responsible for transport of a wide variety of substrates including ions, sugars, nucleobases, vitamins, amines and amino acids, by both symport and antiport. Recent results on the Na^+^ dependence of conformation in the nucleobase transporter Mhp1, which is from a family closely related in structure to LeuT, were markedly different from studies with LeuT (52) portending even greater mechanistic diversity among the many transporter families with the LeuT fold. These observations suggest that a variety of coupling mechanisms may exist even among transporters with closely related structures.

We are struck by the finding that, for both Na^+^ and substrate, interactions that affect the conformation of LeuT involve both scaffold and bundle domains (Fig. 2C and ref. (25)). These results support the idea that ligands affect conformation by binding near the interface between the domains whose movement relative to one another results in transport. Although our results here are relevant for LeuT and probably other transporters in the NSS family, the principle that ligand-induced conformational change involves binding sites at domain interfaces may prove to have wider applicability in transporters.

## Materials and Methods

<u>Mutant construction, expression and binding assays</u> - Mutants of LeuT and SERT were generated using the Stratagene QuickChange protocol as described (25, 53). LeuT mutants were constructed in the WT or Y265C background carrying an His_8_ tag at the C-terminus, whereas SERT mutants were generated in rSERT X5C, a construct in which five endogenous accessible cysteine residues were replaced (54). In addition, Ser277 was replaced with cysteine to allow measurements of cytoplasmic pathway accessibility (24, 38). All mutations were confirmed by DNA sequencing.

LeuT mutants were expressed in *E.coli* as described (25). Cells were disrupted using an Emulsiflex C-3 homogenizer and the membranes were stored at -80°C until further use. Binding measurements in LeuT mutants were made using the SPA technique (36). LeuT was solubilized from frozen *E. coli* membranes in binding buffer (200 mM NaCl containing 20 mM Tris-Cl pH 7.4, 0.1-0.15% n-dodecyl β-D-maltopyranoside (DDM) and 20% glycerol) at a ratio of 240 μg of membrane protein per ml of binding buffer. Solubilized LeuT was bound to SPA beads and washed into the appropriate buffer as described (25). For each mutant, the Na^+^ dependence of saturating (1 μΜ) [^3^H]-leucine binding was first determined, and subsequent binding measurements were performed at the Na^+^ concentration at which leucine binding was half-maximal. Leucine affinity was determined by the ability of unlabeled leucine to displace [^3^H]-leucine (500 nM for Y108F and S256A, 200 nM for I359A, and 5 nM for all other mutants), and alanine affinity was determined similarly.

<u>Reconstitution and Transport assays</u> - Reconstitution of LeuT used a modification of the procedure of Kanner (55). 0.3 g of membrane pellet (wet weight) was solubilized in 6 ml of 20 mM Tris-Cl buffer, pH 8 containing 200 mM NaCl and 40 mM DDM. After centrifugation at 140,000 x g for 40 min, the supernatant fraction was incubated with 600 μ! of nickel affinity resin, Ni-NTA agarose (Qiagen), in the solubilization buffer overnight at 4°C with gentle agitation. The resin was washed 3 times with 20 mM Tris-Cl buffer, pH 8 containing 200 mM NaCl, 4 mM n-decyl β-D-maltopyranoside (DM) and 40 mM imidazole; LeuT was eluted with the same buffer containing 300 mM imidazole and quantified by A_280_ using an extinction coefficient of 1.935 for a 1 mg/ml solution.

The lipid mixture for reconstitution consisted of equal amounts (by weight) of asolectin and *E. coli* polar lipids suspended in 120 mM K-phosphate, pH 7.4 and sonicated until translucent at a concentration of 15 mg/ml. 2 ml of the sonicated lipids was mixed with 1.1 ml of 1.38 M NaCl, and stored at -20°C until use. 175 μl of this lipid mixture was mixed with 10 μl of 20% sodium cholate and then 20 μg of purified LeuT in 20 μl was added for a protein/lipid ratio of 1:100 (w/w). The LeuT-lipid-cholate mixture was incubated for 10 min on ice and applied to a 55 × 4.8 mm column of Sephadex G-50 fine that had been equilibrated in 120 mM Κ-Phosphate, pH 7.4 at 4°C and subjected to pre-centrifugation in a swinging bucket rotor to remove excess liquid but not to dry out the column (approximately 2000 rpm). After application of the lipid-protein mixture, the column was subjected to centrifugation again for 20 s and the eluate, containing reconstituted LeuT proteoliposomes equilibrated with Κ-phosphate buffer, was collected.

To measure [^3^H]alanine transport, 10 μl of reconstituted proteoliposome suspension was diluted into 360 μl of transport buffer (20 mM HEPES-Tris, pH 7 containing 200 mM NaCl) and 40 nM [^3^H]alanine or a control solution in which NMDG-Cl replaced NaCl. After incubation for time intervals of 0-20 min, the reaction was stopped by rapid addition of 3 ml of ice-cold 20 mM HEPES-Tris buffer, pH 7 containing 200 mM NMDG-Cl followed by filtration on 0.45 μm HAWP filters (Millipore). An additional 3 ml of buffer was used to rinse the reaction tube and the filter. Filters were counted in 4 ml of Optifluor (PerkinElmer Life Sciences). Transport rates were calculated from the initial time points during which accumulation approximated a linear time course. For counterflow experiments, the column equilibration buffer was transport buffer containing 5 mM unlabeled alanine. Eluted proteoliposomes were cooled to 4°C, applied to a second Sephadex column equilibrated in transport buffer and subjected to centrifugation at 4°C to remove extravesicular alanine.

Transport of 10 nM [^3^H]proline into bacterial membrane vesicles was performed as described (42) using osmotic shock vesicles (56) and membranes from cells processed in an Avestin C3 cell disrupter. Incubation mixtures consisted of 500 μl of 50 mM Κ-phosphate, pH 6.6, 10 mM MgCl_2_, 50 mM NaCl, 40 μg of membrane proteins, and 20 mM D-lactate. Proline transport was initiated by addition of 10 nM [^3^H]proline (120 Ci/mmol, American Radiolabeled Chemical, St. Louis, MO) and incubation for intervals of 0-10 min at room temperature. The assays were terminated by adding 500 μl of ice-cold 0.1 M LiCl, followed by rapid filtration through HAWP filters (0.45μm, Millipore) and washing with 2 ml of 0.1 M LiCl. Filters were counted in 4 ml of Optifluor.

<u>Cysteine accessibility measurements</u> - Accessibility of the LeuT extracellular or cytoplasmic pathways were determined from the MTSEA (2-aminoethyl methanethiosulfonate) reactivity of LeuT mutants with cysteine residues replacing Val33 or Tyr265, respectively, as described (25). In brief, membranes from *E. coli* expressing a given mutant were trapped on glass-fiber filters and treated with a range of MTSEA concentrations (0.1 μΜ-1 mM unless stated otherwise) for 15 min ± 200 mM Na^+^, or with Na^+^ and alanine. Each sample was then washed from residual MTSEA, denatured and solubilized, and bound to an immobilized Ni^2+^ resin via a C-terminal His_8_ tag. Cysteine residues unmodified by MTSEA were labeled with a fluorescent maleimide, purified and separated by SDS-PAGE. Fluorescence intensity, reflecting the fraction of unmodified cysteine, was quantified, and the MTSEA concentration leading to half-maximal modification determined. This value was converted to a pseudo-first order rate constant (see SI “Modification rates and ratios”). Experimental uncertainties were calculated from the standard deviation associated with the half-maximal MTSEA concentration and the uncertainties in MTSEA concentration and experimental timing with appropriate propagation of uncertainty as the experimental data was processed.

Accessibility measurements with SERT mutants were made as described previously (38). Modification of a cysteine in the cytoplasmic pathway with MTSEA blocks binding of high-affinity ligands by preventing opening of the extracellular pathway. We followed the modification reaction by the decrease in either RTI-55 β-CIT)(38) or citalopram binding (this study).

Data Analysis — Data from binding, transport, and accessibility measurements were fit, plotted, and statistically analyzed using Origin (OriginLab, Northampton, MA).

<u>Molecular simulations</u> - The outward-open conformation of LeuT (PDB 3F48) was used as a starting point for molecular dynamics simulations, including sodium ions, the alanine substrate and the crystallographic waters found in the structure. Residues Glu112, Glu287 and Glu419 were protonated, whereas Glu290 was deprotonated, in line with previous work (25). DOWSER (57) was used to add waters to any unfilled protein cavities. The thus-prepared protein was inserted in a hydrated 233-lipid dimyristoylphosphocholine (DMPC) bilayer, which was previously optimized for the LeuT structures using GRIFFIN (58), as previously described (25). The final system consisted of 39 chloride ions, 33 free sodium ions, and a total of 90,383 atoms, which resulted in a box size of 94 × 94 × 100 Å.

Simulations were performed in the NPT ensemble using the NAMD 2.12 package (59) with the all-atom CHARMM 36 force field (60-63). A constant temperature of 310 Κ was maintained through Langevin dynamics, and a constant pressure of 1 atm was achieved with the Nose’-Hoover Langevin piston algorithm implemented in NAMD. The non-bonded interactions were smoothly switched off from 10 to 12 Å using the van der Waals force switching option, and particle mesh Ewald (PME) summation was used to compute long range electrostatic interactions (64).

The hydrated protein-lipid system was energy-minimized for 100,000 steps using the steepest descent algorithm and the system was subsequently equilibrated in multiple stages. First, harmonic position restraints were progressively reduced on side-chain atoms, backbone heavy atoms and finally Cα atoms over the course of 12 ns. Then, the root-mean-square-deviation (RMSD) of all Cα atoms was restrained to the starting structure, allowing the protein to slightly tumble in the bilayer for 2 ns. Finally, all restraints were released and the protein was allowed to equilibrate for another 10 ns. Subsequent sampling of the wild-type system was considered production. Three production runs were started with a different random seed, each lasting for 700 ns, sampling a total of 2.1 μs.

Simulations of the inward-open conformation of LeuT (PDB 3TT3) were extended to a length of 2 μs from three 700-ns trajectories reported previously (49).

Analysis of trajectories was performed with VMD (65) and MDTraj (66), sampling every 10 or 100 ps.

## Acknowledgments

This work was supported by NIH grants DA007259, DA008213 and NS102277 (to GR) and by the Division of Intramural Research of the NIH, National Institute of Neurological Disorders and Stroke (to LRF). This work utilized the computational resources of the NIH HPC Biowulf cluster (http://hpc.nih.gov). We thank Dr. Titus Boggon for his help in visualizing non-protein densities in the S2 region.

## Abbreviations

NSS, Neurotransmitter:Sodium Symporter; MD, Molecular Dynamics; MTSEA, 2-aminoethyl methanethiosulfonate hydrobromide; NMDG^+^, N-methyl-D-glucamine; FRET, Fӧrster resonance energy transfer; LMNG, lauryl maltose neopentyl glycol; SPA, scintillation proximity assay; SERT, serotonin transporter; NET, norepinephrine transporter; DAT, dopamine transporter; 5-HT, serotonin, 5-hydroxytryptamine; β-CIT, (-)2β-Carbomethoxy3β-(4-iodophenyl)tropane, WT, wild type; DDM, *n*-dodecyl β-D-maltopyranoside; DM, *n*-decyl β-D-maltopyranoside;

## Supporting Information

**Modification rates and ratios**. As described previously (25), modification of Cys265 followed a first-order time course. Rate constants for modification were determined from the concentration of MTSEA that half-maximally modified LeuT mutants in a 15 min incubation under the specified conditions. The half maximal MTSEA concentration was interpolated from extents of modification over a range of MTSEA values (usually 0.1 μΜ to 1 mM). At the half-maximal concentration, the half-time is the assay time - 15 min - and the first-order rate constant is 7.7×10^−4^ sec^−1^. This rate constant was divided by the half maximal MTSEA concentration to obtain the pseudo first-order modification rate constant.

The extent of modification was determined by labeling samples of MTSEA-modified LeuT with a fluorescent maleimide that could be detected in a LiCor Odyssey imager (25). Over a range of experiments using different LeuT mutants, the signal to background ratio for measurements varied from approximately 500 for LeuT unmodified by MTSEA to 10 for maximally-modified LeuT. Figure S8 shows images of the Y265C mutant modified with the same range of MTSEA concentrations ± alanine in the presence of Na^+^, along with plots of the fluorescence values.

**Figure S1.**
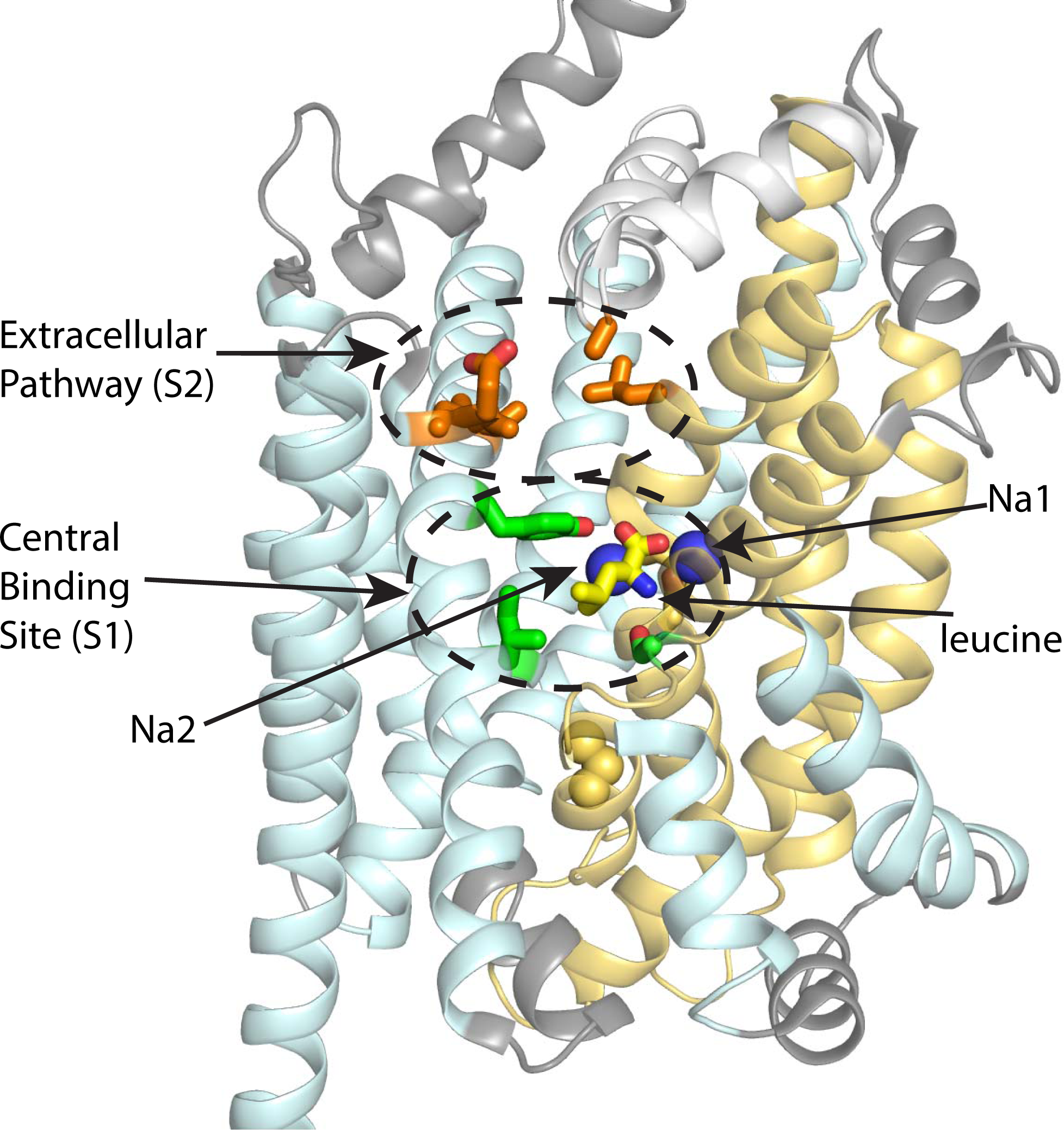
Location of the alanine binding site (lower dashed oval) in the overall structure of outward-occluded LeuT (snapshot after ~500 ns simulation of PDB 3F48). The protein is shown as cartoon helices, with the scaffold domain (TMs 3–5 and 8–12) and the bundle domain (TMs 1, 2, 6 and 7) colored pale cyan and yellow-orange, respectively. Long extracellular and intracellular loops are colored gray except for EL4 (white). Binding site residues are in stick representation, with mutated substrate binding site (S1) residues in green while Na^+^ and residue 265 are shown as spheres. TM helix 11 is omitted for clarity. The upper dashed oval shows the location of the proposed S2 site in the overall structure (snapshot after 2 μs simulation based on PDB 3TT3 (49)). Mutated residues in the region of S2 are colored orange.

**Figure S2.**
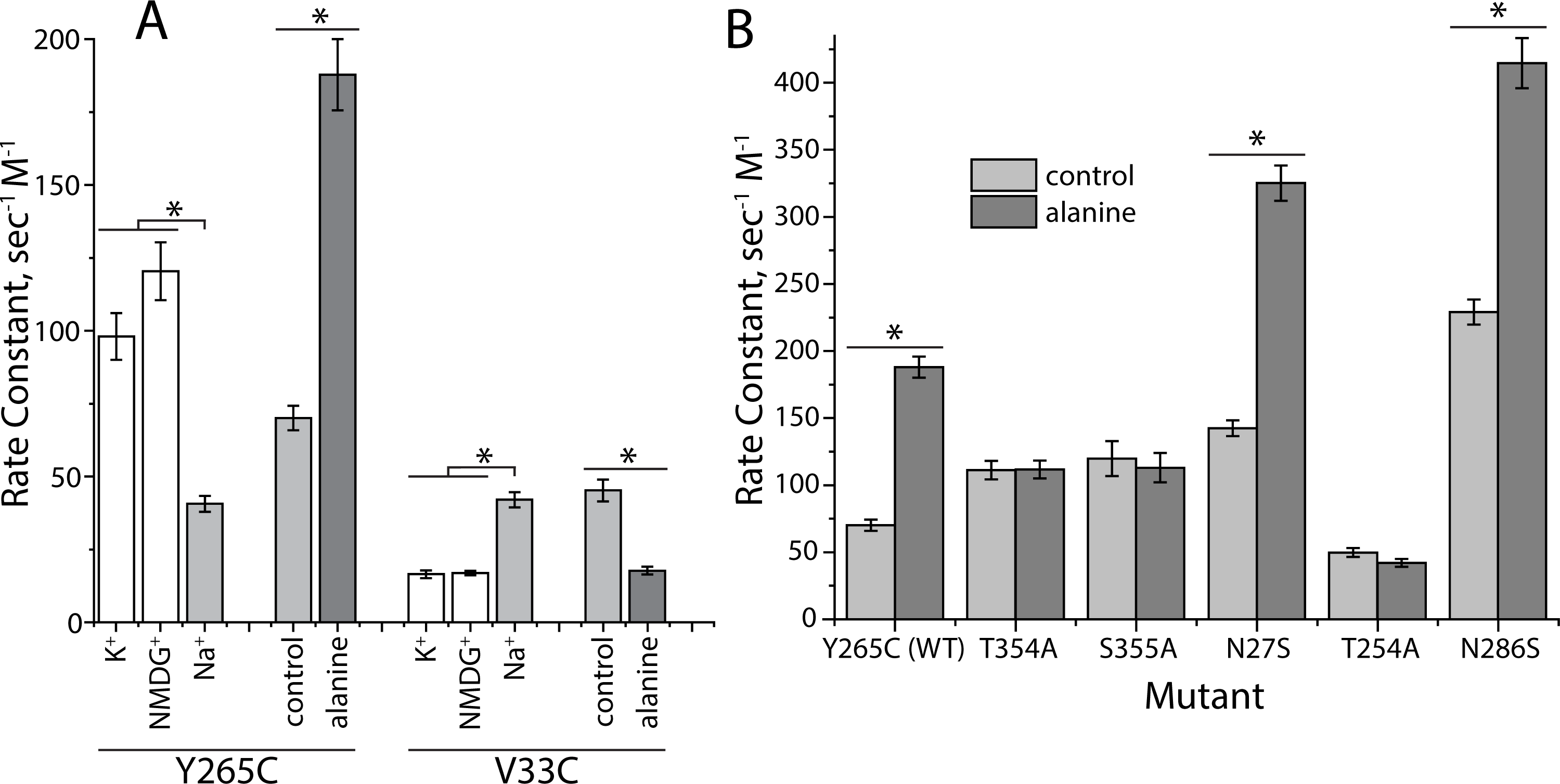
*Individual Rate Constants.* A. Effect of Na^+^ and alanine on rate constants for MTSEA modification of V33C (extracellular pathway) and Y265C (cytoplasmic pathway). For Y265C, n=5 for ± Na^+^, n=3 for ± alanine. Both alanine and control rates were measured in Na^+^. For V33C, n=5 for ± Na^+^, n=4 for ± alanine. B. Effect of Na1-site mutations N27S, T254A and N286S and Na2-site mutations T354A and S355A on the rate constants for Y265C modification in Na^+^ alone and Na^+^ plus alanine. Asterisks indicate a significant difference (P < 0.05 in paired t-test) resulting from alanine addition. For Y265C and N286S n=4, n=3 for T354A, N27S and T254A, n=2 for S355A.

**Figure S3.**
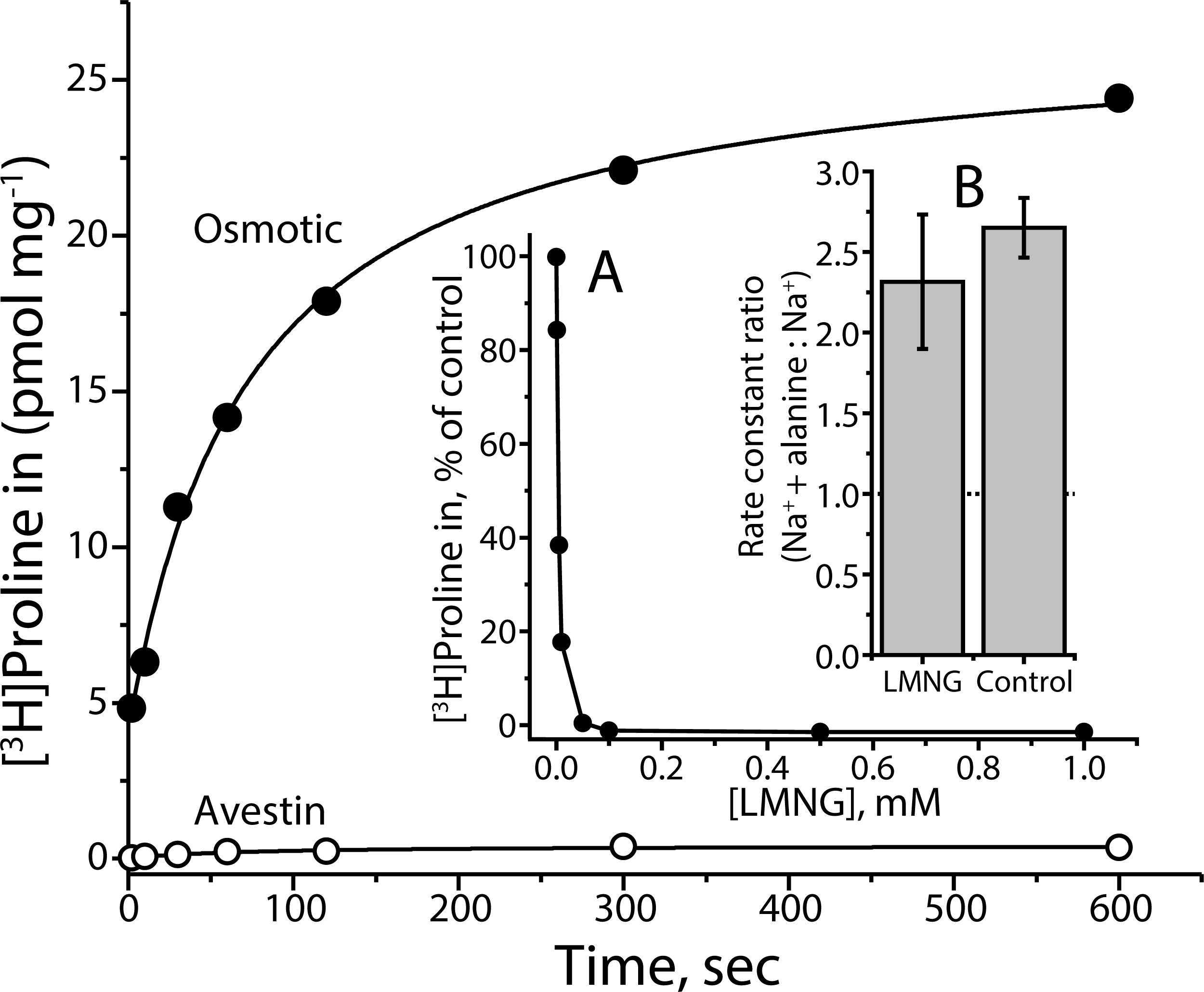
*Substrate-dependent conformational changes in LeuT do not require a transmembrane concentration gradient.* Membranes were prepared from *E. coli* expressing LeuT Y265C using the osmotic shock procedure (filled circles) (56) or an Avestin C3 cell disrupter (open circles) and influx of 10 nM [^3^H]proline was measured in the presence of 20 mM D-lactate, which is a substrate for generation of electrochemical H+ and Na^+^ potentials in *E. coli.* After the indicated incubation time, membranes were filtered, washed and counted as described (56); representative data from one of 2 experiments. *Inset* A. The indicated concentrations of the detergent LMNG were added to membranes, and incubated for 5 min with [^3^H]proline; representative data from one of 2 experiments. Separate experiments using the amido black assay for filter-bound protein (67) demonstrated that, up to 1 mM LMNG, the amount of membrane protein trapped on the filters was unchanged. *Inset* B. Rate constant ratio for 20 μM alanine + 10 mM Na^+^ relative to Na^+^ alone in the presence or absence of 0.2 mM LMNG. There was no significant difference with or without LMNG detergent; n=2.

**Figure S4.**
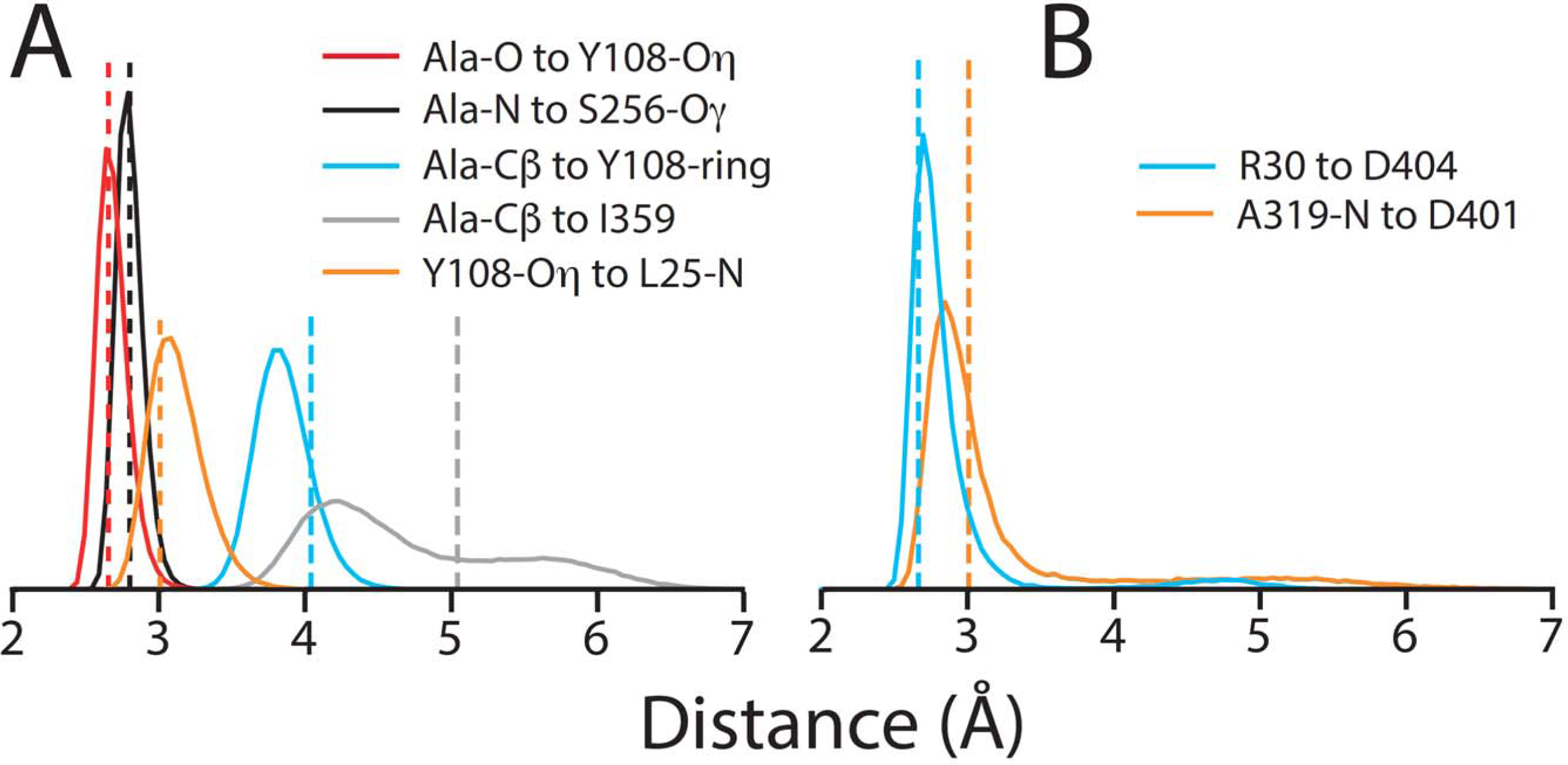
*Relevant interactions observed in the detergent-solubilized crystal structures of LeuT are maintained during molecular dynamics simulations.* Distance distributions for key interacting groups were computed for molecular simulations of LeuT in a lipid bilayer (solid lines) relative to the values in the starting crystal structures (dashed lines) for two states of the transporter; representative frames of those trajectories are shown in Fig. 2A and 3A. (A) Simulations of outward-occluded alanine-bound LeuT (PDB 3F48), analyzed every 10 ps over three 700 ns-long trajectories. Distribution of distances between substrate atoms and the side-chains of key residues in the S1 binding site. Distances to the Tyr108 ring and to Ile359 were measured to the nearest carbon atom. (B) Simulations of substrate-free inward-open LeuT (PDB 3TT3) analyzed every 100 ps over three 2µs trajectories, which were extended from the 700ns-simulations of Adhikary et al. (49). Despite its hydrated location close to the surface, the hydrogen-bond between the backbone nitrogen of Ala319 in extracellular loop 4 and the nearest side chain carboxyl oxygen of Asp401 is preserved, with only rare forays to distances of 4–6 Å (orange). Similar features are found in the closest distance between Arg30 terminal nitrogen and Asp404 carboxyl oxygen atoms (cyan).

**Figure S5.**
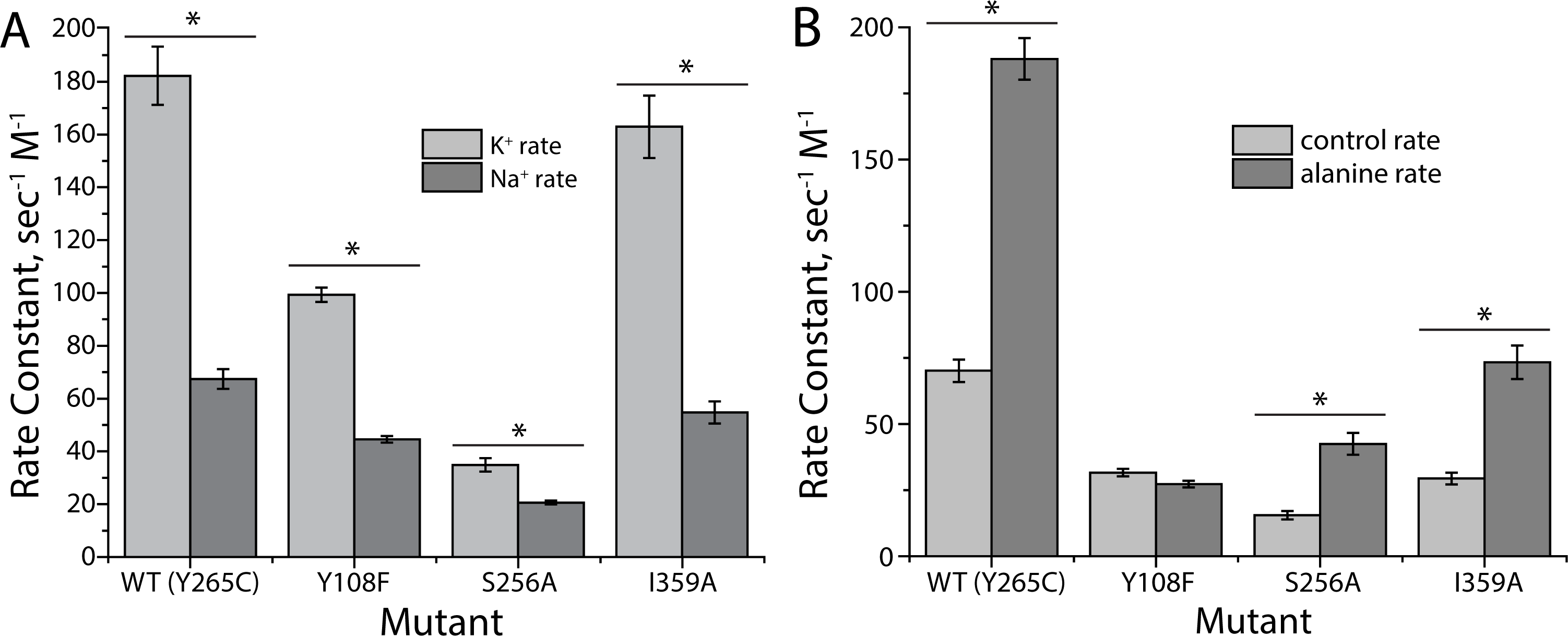
*Conformational changes in substrate binding site mutants.* **A.** Effect of Na^+^ on rate constants for MTSEA modification of Y265C (control), Y108F, S256A and I359A (all in the Y265C background). Aside from the decreased accessibility in the presence of Na^+^ for all mutants, there was an overall decrease in reactivity in these mutants consistent with a shift toward inward-closed conformations. **B.** Effect of alanine in the presence of Na^+^, relative to Na^+^ alone, on rate constants. Aside from the effect of mutation on absolute rates noted above, an increase in Cys265 accessibility in the presence of alanine was observed for Y265C, and S256A and I359A (in the Y265C background). No increase was observed in Y108F. Asterisks indicate a significant difference (P < 0.05 in paired t-test) resulting from Na^+^ (A) or alanine (B) addition; n=4 except S256A ± alanine, n=3.

**Figure S6.**
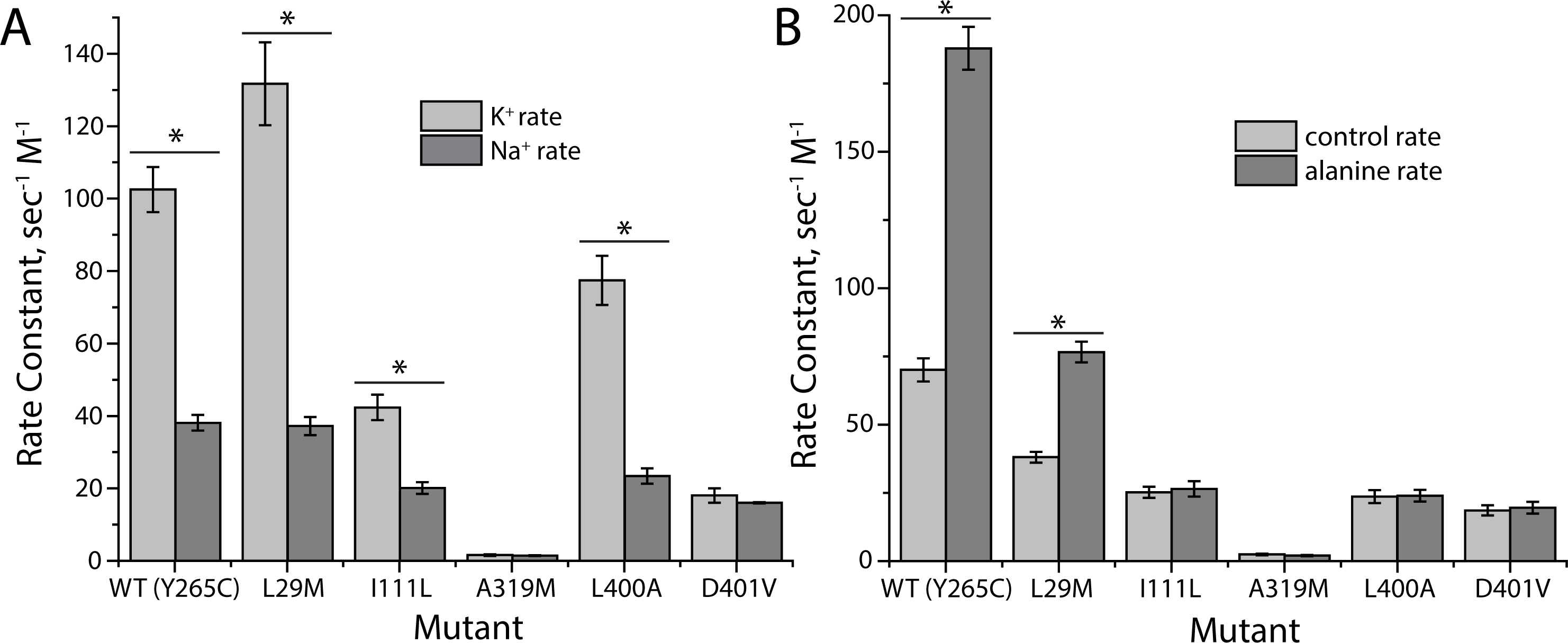
Conformational changes in extracellular pathway mutants. Effect of Na^+^ (A) and alanine (B) on rate constants for MTSEA modification of Y265C (control), L29M, I111L, A319M, L400A and D401V. Control rates (K+) for MTSEA modification of Cys265 were moderately suppressed for I111L and D401V, but dramatically suppressed for A319M (also in B), suggesting that in this mutant, the extracellular pathway is unable to close. Control rates in B included Na^+^. Asterisks indicate a significant difference (P < 0.05 in paired t-test) resulting from Na^+^ (A) or alanine (B) addition. Replicates are as in Figure 3B.

**Figure S7.**
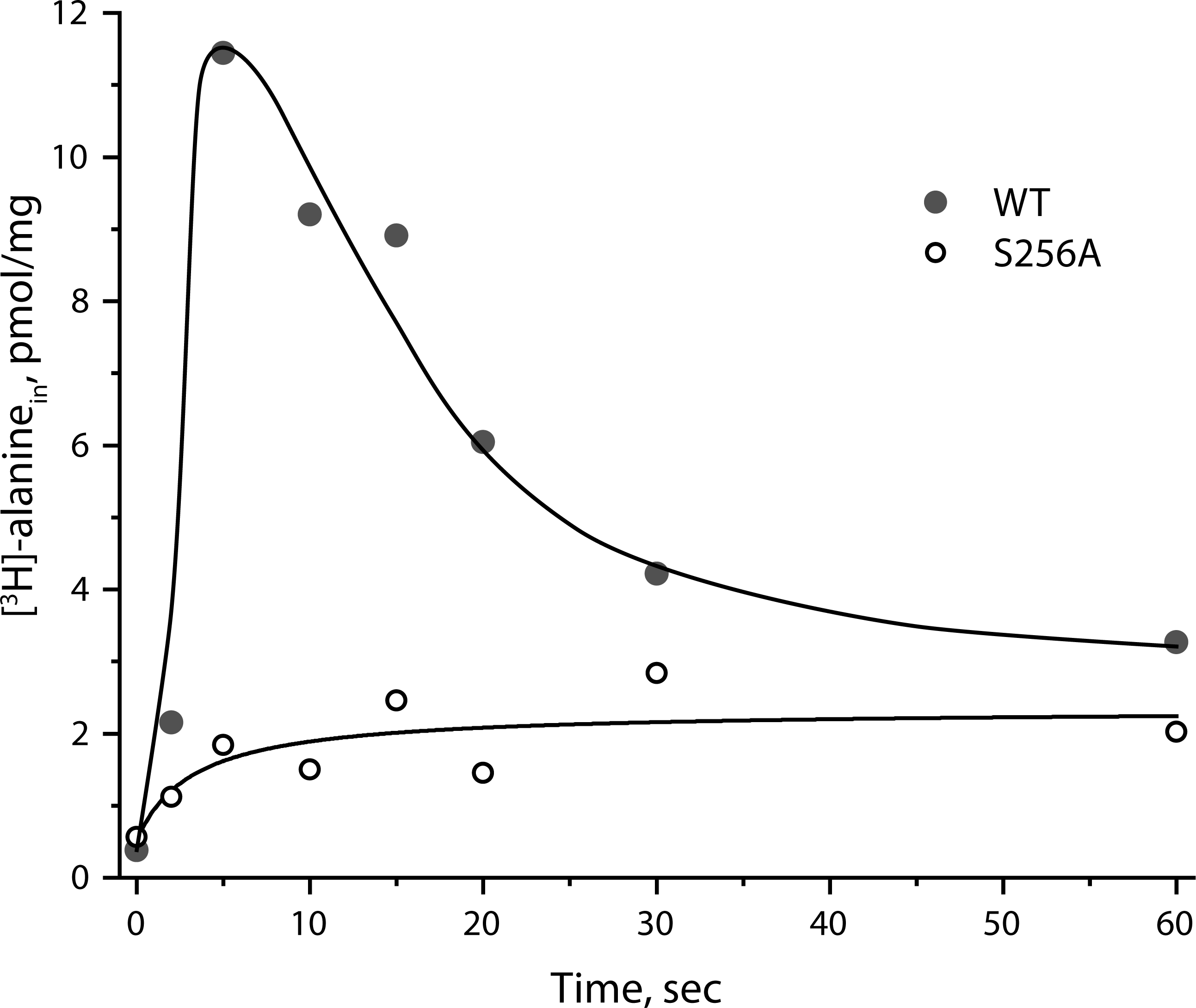
*Wild type but not S256A LeuT catalyzes substrate counterflow.* Proteoliposomes incorporating either LeuT WT or S256A and containing transport buffer with 5 mM unlabeled alanine were prepared as described under “Methods”. 40-fold dilution of WT liposomes into transport buffer containing [^3^H]-alanine led to a transient accumulation of radiolabel followed by a decline as unlabeled alanine exited the liposomes. Proteoliposomes incorporating S256A LeuT did not accumulate radiolabel transiently. Displayed values represent the accumulation above a control in which unlabeled alanine was omitted from the loading solution. The experiment shown was performed a total of 3 times with similar results.

**Figure S8.**
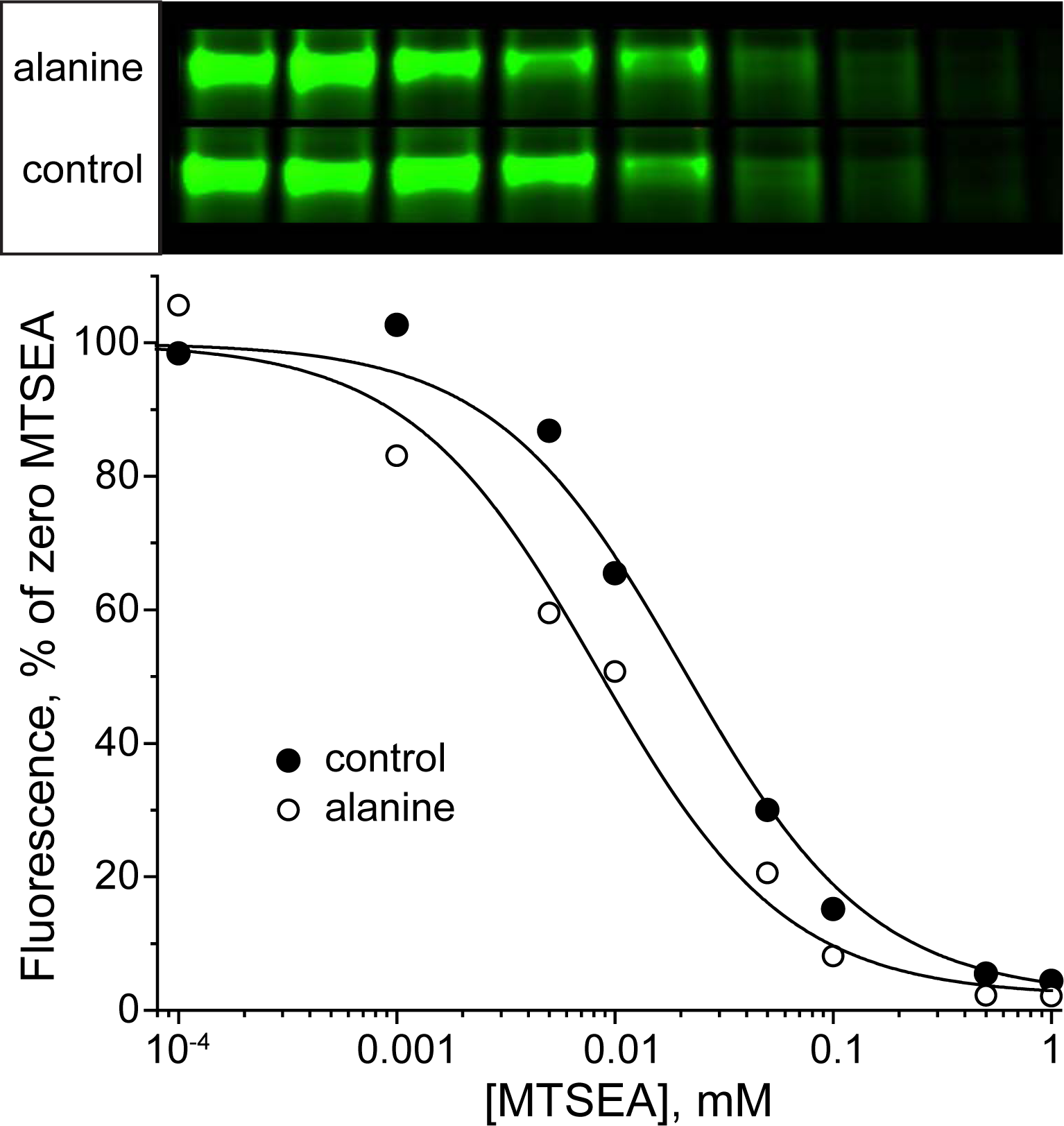
*Example of primary data used for calculation of rates and ratios.* From an experiment comparing the accessibility of Cys265 in the presence of Na^+^ ± alanine, the top panel shows the fluorescent LeuT bands from samples with and without alanine present during treatment with MTSEA. The lower panel shows a plot of the fluorescence intensities vs. MTSEA concentration. In this experiment, half-maximal modification occurred at 8.3 μM MTSEA in the presence of alanine and 20.6 μM in the control, yielding pseudo first-order rate constants of 84 and 34 sec^−1^ M^−1^, respectively and a ratio of 2.47. Typically, 3–5 experiments of this type were used to calculate the Na^+^ or alanine ratio for each mutant.

